# Glucosinolate and phenylpropanoid biosynthesis are linked by proteasome-dependent degradation of PAL

**DOI:** 10.1101/668038

**Authors:** Jeong Im Kim, Xuebin Zhang, Pete E. Pascuzzi, Chang-Jun Liu, Clint Chapple

## Abstract

Plants produce several hundreds of thousands of secondary metabolites that are important for adaptation to various environmental conditions. Although different groups of secondary metabolites are synthesized through unique biosynthetic pathways, plants must orchestrate their production simultaneously.

Phenylpropanoids and glucosinolates are two classes of secondary metabolites that are synthesized through apparently independent biosynthetic pathways. Genetic evidence has revealed that the accumulation of glucosinolate intermediates limits phenylpropanoid production in a Mediator Subunit 5 (MED5) dependent manner.

To elucidate the molecular mechanism underlying this process, we analyzed the transcriptomes of a suite of glucosinolate-deficient mutants using RNAseq and identified mis-regulated genes that are rescued by the disruption of MED5. The expression of a group of *Kelch Domain F-Box* genes (*KFB*s) that function in PAL degradation is affected in glucosinolate biosynthesis mutants and the disruption of these *KFBs* restores phenylpropanoid deficiency, dwarfism and sterility in the mutants.

Our study suggests that glucosinolate/phenylpropanoid metabolic crosstalk involves the transcriptional regulation of KFB genes that initiate the degradation of the enzyme phenylalanine ammonia-lyase, which catalyzes the first step of the phenylpropanoid biosynthesis pathway. Nevertheless, KFB mutant plants remain partially sensitive to glucosinolate pathway mutations, suggesting that other mechanisms that link the two pathways also exist.

## Introduction

Phenylpropanoids are a group of metabolites made primarily from the amino acid phenylalanine and include a variety of hydroxycinnamoyl derivatives, anthocyanins and flavonols, as well as other low molecular weight conjugates. Another product of phenylpropanoid metabolism is lignin, a major component of the cell wall that provides rigidity and cell wall hydrophobicity to the vasculature. Plants belonging to Brassicales produce glucosinolates, which function as defense molecules by deterring herbivores and pathogens. Over 100 different types of glucosinolates have been found in plants (Halkier, 2016; Agerbirk & Olsen, 2012). Glucosinolates that are derived from tyrosine and phenylalanine are called aromatic glucosinolates whereas glucosinolates derived from tryptophan or from aliphatic amino acids are called indole or aliphatic glucosinolates, respectively (Halkier & Gershenzon, 2006). It has been shown that both the phenylpropanoid and glucosinolate biosynthetic pathways can be regulated at various levels including the transcriptional, and post-translational levels, as well as at the metabolic level through feed-forward and feed-back enzyme inhibition (Yin *et al.*, 2012; Nintemann *et al.*, 2017). In addition to the regulation of their own pathways, there is crosstalk between the glucosinolate and phenylpropanoid pathways (Hemm *et al.*, 2003; Kim *et al.*, 2015). Studies with two *Arabidopsis* mutants, *<u>r</u>educed <u>e</u>pidermal <u>f</u>luorescence2* (*ref2*) and *ref5*, demonstrated that the accumulation of glucosinolate biosynthetic intermediates can limit the production of phenylpropanoids (Hemm *et al.*, 2003; Kim *et al.*, 2015). The *ref5* and *ref2* mutants were originally isolated from a screen that was aimed at the isolation of phenylpropanoid-deficient mutants (Ruegger & Chapple, 2001). Wild-type *Arabidopsis* plants fluoresce blue upon ultra violet (UV) light exposure due to the accumulation of sinapoylmalate, a UV-protective phenylpropanoid in the leaf epidermis (Ruegger & Chapple, 2001; Landry *et al.*, 1995). Plants having defects in sinapoylmalate accumulation fluoresce red because the absence of sinapoylmalate enhances and reveals UV-induced chlorophyll fluorescence in the mesophyll cells. Using this characteristic color difference, a series of *ref* mutants were isolated, and their characterization led to the identification of several key enzymes and regulators of phenylpropanoid biosynthesis (Meyer *et al.*, 1996; Ruegger & Chapple, 2001; Franke *et al.*, 2002; Stout *et al.*, 2008; Schilmiller *et al.*, 2009; Bonawitz *et al.*, 2012). Unexpectedly, genetic mapping studies revealed that *ref2* and *ref5* mutants have mutations in *CYP83A1* (*REF2*) (Hemm *et al.*, 2003), and *CYP83B1* (*REF5*) (Kim *et al.*, 2015), which are involved in aliphatic and indole glucosinolate biosynthesis, respectively. Consistent with the biochemical functions of CYP83A1 and CYP83B1 in the synthesis of aliphatic and indolic glucosinolates, *ref2* and *ref5* mutants also show glucosinolate deficiency phenotypes, but the absence of these compounds does not lead to alterations in phenylpropanoid metabolism. Instead, biochemical and genetic analyses revealed that the accumulation of aldoximes or their derivatives negatively impacts the phenylpropanoid pathway (Kim *et al.*, 2015).

When the production of indole-3-acetaldoxime (IAOx), the substrate of REF5/CYP83B1, was blocked by knocking out the two enzymes that are responsible for converting tryptophan to IAOx, CYP79B2 and CYP79B3, indole glucosinolates were eliminated and the phenylpropanoid content of *ref5* was increased (Kim *et al.*, 2015). These data suggest that IAOx, or a derivative thereof, is responsible for the crosstalk between the glucosinolate and phenylpropanoid pathways. In agreement with this model, increased production of IAOx induced by overexpression of *CYP79B2* in wild type represses the production of phenylpropanoids (Kim *et al.*, 2015). Importantly, *cyp79b2 cyp79b3* double mutants, which lack IAOx, have elevated levels of soluble phenylpropanoids, indicating that this cross-pathway interaction occurs in wild-type plants.

A *ref5* suppressor screen revealed that MED5a, a subunit of the transcriptional co-regulator Mediator complex, is involved in the crosstalk between glucosinolate and phenylpropanoid metabolism. *Arabidopsis* has two MED5 paralogs, MED5a and MED5b (REF4)(Bonawitz *et al.*, 2012, 2014). *med5a/b* double mutants contain substantially increased levels of phenylpropanoids and exhibit increased expression of several phenylpropanoid biosynthesis genes whereas *med5a* and *med5b* single mutants exhibit either no obvious phenotype or a more modest phenotype compared to *med5a/b* (Bonawitz *et al.*, 2012, 2014). Although the accumulation of aldoximes or their derivatives represses the early step(s) of phenylpropanoid pathway through MED5, it is still not clear how aldoxime metabolism affects the phenylpropanoid pathway.

To understand the molecular mechanisms underlying the crosstalk, we conducted transcriptome profiling using a suite of glucosinolate-deficient mutants. Here we report that RNAseq analysis identified 71 candidate genes that are mis-regulated in *ref5* and *ref2* in a MED5-dependent manner. We show that the expression of a set of F-box genes functioning in PAL ubiquitination is increased in both *ref5* and *ref2*. Consistent with the known function of these F-box proteins, PAL activity and the amount of PAL protein are reduced in *ref5* and *ref2* and disruption of these F-box genes restores the phenylpropanoid deficiency in the mutants.

## Materials and Methods

### Plant Material and Growth Conditions

*Arabidopsis thaliana* (ecotype Col-0) plants were cultivated at a light intensity of 100 μE m^−2^ sec^−1^ at 22°C under a photoperiod of 16 hr light/8 hr dark. Seeds were held for 2 days at 4°C before transferring to the growth chamber. The *ref5-1* allele (*ref5*) has a missense mutation in CYP83B1 (Kim *et al.*, 2015), the *ref2-1* allele (*ref2*) has a nonsense mutation in CYP83A1 (Hemm *et al.*, 2003) and the *med5* mutant is *med5a med5b* double mutant which are T-DNA insertion lines (Bonawitz *et al.*, 2012; Kim *et al.*, 2015). The *kfb1-1*, *kfb20-1* and *kfb50-1* alleles were previously reported (Zhang *et al.*, 2013). The *kfb39* mutant and the *myb4* mutant were generated using CRISPR/Cas9 system (see below for detail information) (Panda *et al.*, unpublished). The homozygosity of all mutants used in this study was confirmed by following their genotyping methods as were previously reported (Panda C Wager A Chen H-Y Xu Li; Hemm *et al.*, 2003; Bonawitz *et al.*, 2012; Zhang *et al.*, 2013; Kim *et al.*, 2015). The homozygosity of *kfb39* mutant was confirmed by the presence or absence of *DdeI* site in the CRISPR/Cas9 target sequences (see below for detail information).

### Metabolite analysis

For soluble metabolite analyses, leaves from three-week old plants were extracted with 50% methanol (v/v) at 65°C for 90 minutes at a tissue concentration of 100 mg mL ^−1^. Samples were centrifuged at 16000xg for minutes. 10 μL of extraction was loaded on a Shim-pack XR-ODS column (Shimadzu, Japan) and separated at an increasing concentration of acetonitrile from 2% to 25% for 18 min in 0.1% formic acid at a flow rate of 0.8 mL min^−1^. Compounds were identified based on their retention times and UV spectra. Sinapoylmalate was quantified using sinapic acid as a standard.

### High-throughput mRNA sequencing

For RNAseq analysis, we collected three biological replicates of whole rosette leaves from 18-day-old plants. Total RNA was prepared using the Qiagen RNeasy Plant Mini Kit (Qiagen Sciences) following the manufacturere's instructions. RNA samples were treated with RNase-free DNase Set (Qiagen Sciences) according to the manufacturer's instructions. Total RNA was submitted to the Purdue Genomics Core Facility (Purdue University) for library construction and sequencing. cDNA libraries for sequencing were prepared using a TruSeq Stranded mRNA Library Prep Kit (Illumina) according to the manufacturer’s instructions but 8 cycles of PCR amplification were performed rather than the recommended 15. All 18 libraries were then titrated using a KAPA quantification kit (Kapa Biosystems) and added to a single library pool. That pool was denatured and clustered on two lanes of Illumina High Output Chemistry flow cells at 16.5 pM. Paired-end, 100-bp sequencing was performed by an Illumina HiSeq 2500 in rapid mode. To pre-process filtered Illumina reads for mapping, sequences were first assessed for quality using FastQC (v.0.11.2; http://www.bioinformatics.babraham.ac.uk), then trimmed using the Trimmomatic toolkit (v.0.32; http://www.usadellab.org/cms/?page=trimmomatic) to remove bases with a Phred33 score less than 20 and a minimum length of 30. Quality-trimmed reads were then mapped to the bowtie2- indexed *Arabidopsis* genome (Bowtie ver 2.2.5.0 and TAIR10) using Tophat (v.2.0.13; http://tophat.cbcb.umd.edu/) with default parameters (Trapnell *et al.*, 2009). A counts matrix consisting of the raw read count was generated using HTSeq (v.0.6.1; http://www-huber.embl.de/users/anders/HTSeq/) (Anders *et al.*, 2015). To eliminate fold changes of infinity and divide-by-zero errors, the counts matrix was modified such that genes with 0 counts across all samples were removed, and remaining 0 count values were changed to 1. Differential expression analysis was performed using the statistical program R (v.3.2.0; http://www.r-project.org/) in conjunction with three analytical methods available from Bioconductor (http://www.bioconductor.org): DESeq2 (v.1.8.1), edgeR (v.3.10.2) and the Cufflinks (v.2.2.1) suite of programs (http://cufflinks.cbcb.umd.edu/). All three statistical methods gave similar overall conclusions. We selected the most conservative results (edgeR; false discovery rate, 0.05) for further investigation and reporting (Robinson *et al.*, 2010). Venn diagrams were adapted from those created with the online tool Venny (v.2.0.2; http://bioinfogp.cnb.csic.es/tools/venny/) and gene ontology term analysis was performed using the online tool DAVID (v.6.7; http://david.ncifcrf.gov)(Huang *et al.*, 2009b; Huang *et al.*, 2009a). The RNAseq data used in this study have been deposited in NCBI’s Gene Expression Omnibus (Edgar *et al.*, 2002) under GEO Series accession number GSE99581.

### Generation of *kfb39* knock-out mutant using CRISPR/Cas9 system

To generate *kfb39* mutants, we used the psgRNA-Cas9-KFB39 vector which was previously reported to generate *Arabidopsis* mutants (Feng *et al.*, 2014). The psgRNA-CAS9 vector was cut with *Bbs*I and then a *KFB39* target sequence was inserted into a cut psgRNA-CAS9 vector using the primer pair CC4841 (5’- GATTGCA TCG CGT TGC GTT TCC AG-3’) and CC4842 (5’- AAACCT GGA AAC GCA ACG CGA TGC-3’). The resulting *KFB39* target sequence together with guide RNA and CAS9 was subsequently cloned into pCAMBIA1300 vector after cutting with *Hin*dIII and *Kpn*I. The pCAMBIA1300 vector carrying KFB39-CRISPR construct was introduced into *ref5* and *ref2* plants via *Agrobacterium tumefaciens* (strain C58 pGV3850) mediated transformation. To select hygromycin resistant transformants, T1 seeds were planted on MS media containing hygromycin after surface sterilization. Hygromycin-resistant T1 transgenic *Arabidopsis* plants were transferred to soil and genomic DNA was extracted from leaves to identify mutations in *KFB39*. The target regions were amplified by PCR with the primer pair CC4964 (5’- TGC TCA ATC AAA CAC ACT CTC T-3’) and CC4928 (5’- GTT TCC CAT CGA CGT TCA CT -3’) and the PCR products were sequenced to identify mutations. We found homozygous mutants carrying ‘T’ insertion at the 3 bases upstream of PAM site which causes frame shift resulting in truncated KFB39 protein with 268 amino acids instead of 410 amino acids. Homozygosity of *kfb39* mutation in *ref5 kfb39* and *ref2 kfb39* was confirmed with *Dde*I digestion because the ‘T’ insertion generates a *D*de*I* recognition site.

### PAL activity and PAL protein quantification

For crude protein extraction, 3^rd^ and 4^th^ rosette leaves from 3-week old seedlings were harvested and ground in liquid nitrogen. 500 μL of extraction buffer containing 100 mM Tris-HCl pH 8.3, 2 mM dithiothreitol, and 10% (v/v) glycerol was added and the mixture was stirred at 4°C for 1 hour. After centrifugation at 10,000xg for 10 min at 4°C, the supernatant was collected, desalted into extraction buffer on a column filled with Sephadex G-50 (GE Healthcare Life Science, Pittsburgh, USA) and assayed for PAL activity. Total protein content was measured using Quick Start Bradford protein assay kit (Bio-Rad, Hercules, CA USA) using bovine serum albumin as a standard. To measure the activity of PAL, 100 μL aliquots of the protein extracts were incubated with 4 mM Phe in reaction buffer (100 mM Tris-HCl pH 8.3, 2 mM DTT, 10% (v/v) glycerol) in a total volume of 280 μL. Reactions were incubated at 37°C for 60 min and then stopped by addition of 20 μL of glacial acetic acid. Reaction products were extracted with 400 μL ethyl acetate, 300 μL of which was removed, dried in a speed vac and redissolved in 50 μL of 50% methanol. 10 μL of the final extract was analyzed by HPLC to quantify the reaction product, cinnamic acid. The PAL activity and protein quantification shown in Figure 5a were performed with whole rosette leaves from three-week old plants by following previous reports (Zhang *et al.*, 2015).

## Results

### Disruption of MED5 rescues the phenylpropanoid deficiency in *ref5* and *ref2* mutants

As shown in Fig. 1a and b, *ref5* and *ref2* mutants appear reddish under UV, whereas wild-type plants fluoresce blue due to the accumulation of sinapoylmalate. Consistent with their *ref* phenotype, both *ref5* and *ref2* mutants contain reduced levels of sinapoylmalate compared to wild type (Fig. 1b). *ref5 med5* (*ref5 med5a/b*) restores the phenylpropanoid accumulation in *ref5* as was previously reported (Fig. 1a, b) (Kim *et al.*, 2015). Because CYP83A1/REF2 and CYP83B1/REF5 functions similarly in the glucosinolate biosynthetic pathway (Hemm *et al.*, 2003; Kim *et al.*, 2015), we speculated that the disruption of MED5 would rescue the phenylpropanoid deficiency of the *ref2* mutant. To test this hypothesis, *ref2 med5* (*ref2 med5a/b*) plants were generated and their biochemical phenotype was assessed. The *ref2 med5* mutant shows bluish fluorescence under UV and the level of sinapoylmalate in *ref2 med5* is increased compared to the *ref2* single mutant, indicating an involvement of MED5 in the crosstalk in *ref2* (Fig. 1a, b). It is important to note, however, that the sinapoylmalate content of *ref5 med5* and *ref2 med5* is less than that of *med5*, suggesting that there may be MED5-dependent and -independent mechanisms involved in glucosinolate/phenylpropanoid crosstalk. Alternatively, given the repressive effect of MED5 on the phenylpropanoid pathway, we continued to consider the possibility that the biochemical phenotype of the *med5* mutations in some way bypasses the phenylpropanoid deficiency of *ref5* and *ref2* mutants and that MED5 is not actually involved in this interaction. Nevertheless, because the disruption of MED5 restores the phenylpropanoid deficiency in both *ref5* and *ref2* mutants, we hypothesized that there is a common mechanistic basis for how glucosinolate metabolites affect phenylpropanoid production that involves MED5 (Fig. 1c).

**Figure 1.**
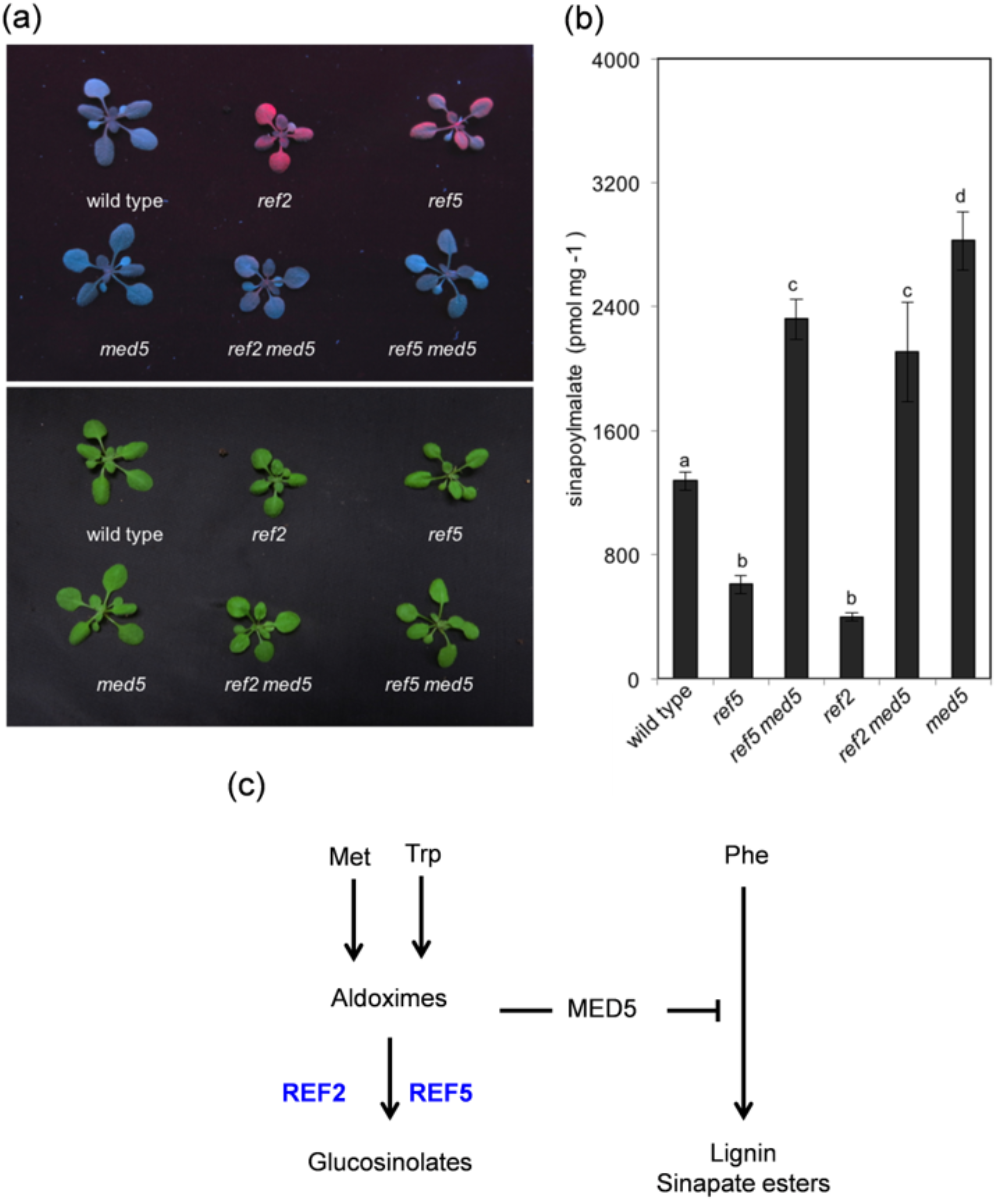
The disruption of MED5 rescues the phenylpropanoid deficiency in *ref5* and *ref2* mutants (a) Representative photographs of wild type (Col-0), *ref2*, *ref5, med5, ref2 med5*, and *ref5 med5* under UV light (top) or white light (bottom). (b) Quantification of sinapoylmalate content in leaves of wild type (Col-0), *ref2*, *ref5, med5, ref2 med5*, and *ref5 med5.* Rosette leaves from three-week-old plants were analyzed for sinapoylmalate content. Data represent mean ± SD (n=3). The means were compared by one-way ANOVA, and statistically significant differences (*p* < 0.05) were identified by Tukey’s test and indicated by a to d to represent difference between groups. (C) Scheme of the crosstalk between glucosinolate biosynthesis and phenylpropanoid biosynthesis. REF2 and REF5, members of the CYP83 family, oxidize aldoximes to products that are precursors of glucosinolates. The accumulation of aldoxime or aldoxime derivatives in *ref5* and *ref2* limits phenylpropanoid production in MED5 dependent way.

### Transcriptome profiles identify mis-regulated genes in glucosinolate-deficient mutants

Given that MED5 is a subunit of a transcriptional co-regulator, it seemed likely that it functions in the crosstalk by influencing the expression of genes that play roles in phenylpropanoid metabolism. To explore this possibility, we performed RNAseq analysis using total RNA prepared from whole rosette leaves of wild type (Col-0), *ref2, ref5, med5, ref2 med5*, and *ref5 med5* to identify genes that are mis-regulated in *ref5* and *ref2* but not in *ref5 med5* and *ref2 med5.* RNAseq analysis identified that 1074 genes were up-regulated in *ref5* compared to wild type and 2423 genes were up-regulated in *ref2* compared to wild type. Among these, 402 genes were increased in both *ref5* and *ref2* mutants compared to wild type (Fig. 2a), 53 of which show restored expression in the absence of MED5 (Fig. 2a, 2c, Table 1). Similarly, we found that 18 genes are down-regulated in both *ref5* and *ref2* compared to wild type in a MED5-dependent manner (Fig. 2b, 2d and Table 2). Thus, this analysis identified a total of 71 candidate genes that may be involved in the MED5-dependent crosstalk between glucosinolate and phenylpropanoid biosynthesis (Table 1, 2).

**Table 1.** The list of 53 genes of which increased expression in *ref2* and *ref5* are restored by the disruption of MED5.

| Gene ID | Name | Description |
| --- | --- | --- |
| AT3G59930 | AT3G59930 | Unknown |
| AT2G44130 | KFB39 | Galactose oxidase/kelch repeat superfamily protein |
| AT3G13310 | AT3G13310 | Chaperone DnaJ-domain superfamily protein |
| AT2G16720 | MYB7 | MYB domain protein 7 |
| AT2G15310 | ARFB1A | ADP-ribosylation factor B1A |
| AT4G16870 | AT4G16870 | transposable element gene |
| AT1G21550 | AT1G21550 | Calcium-binding EF-hand family protein |
| AT2G16890 | AT2G16890 | UDP-Glycosyltransferase superfamily protein |
| AT3G04000 | AT3G04000 | NAD(P)-binding Rossmann-fold superfamily protein |
| AT4G14690 | ELIP2 | Chlorophyll A-B binding family protein |
| AT1G10140 | AT1G10140 | Uncharacterized conserved protein UCP031279 |
| AT1G51090 | AT1G51090 | Heavy metal transport/detoxification superfamily protein |
| AT1G17830 | AT1G17830 | Protein of unknown function (DUF789) |
| AT4G37400 | CYP81F3 | cytochrome P450, family 81, subfamily F, polypeptide 3 |
| AT4G38620 | MYB4 | myb domain protein 4 |
| AT1G71030 | MYBL2 | MYB-like 2 |
| AT3G59940 | KFB50 | Galactose oxidase/kelch repeat superfamily protein |
| AT4G36040 | J11 | Chaperone DnaJ-domain superfamily protein |
| AT1G22570 | AT1G22570 | Major facilitator superfamily protein |
| AT4G31870 | GPX7 | glutathione peroxidase 7 |
| AT3G22060 | AT3G22060 | Receptor-like protein kinase-related family protein |
| AT5G24160 | SQE6 | squalene monooxygenase 6 |
| AT5G43860 | CLH2 | chlorophyllase 2 |
| AT3G22840 | ELIP1 | Chlorophyll A-B binding family protein |
| AT1G19230 | AT1G19230 | Riboflavin synthase-like superfamily protein |
| AT1G15510 | ECB2 | Tetratricopeptide repeat (TPR)-like superfamily protein |
| AT4G27820 | BGLU9 | beta glucosidase 9 |
| AT1G67360 | AT1G67360 | Rubber elongation factor protein (REF) |
| AT3G30460 | AT3G30460 | RING/U-box superfamily protein |
| AT5G61350 | AT5G61350 | Protein kinase superfamily protein |
| AT3G26290 | CYP71B26 | cytochrome P450, family 71, subfamily B, polypeptide 26 |
| AT4G25640 | DTX35 | detoxifying efflux carrier 35 |
| AT4G39090 | RD19 | Papain family cysteine protease |
| AT3G26740 | CCL | CCR-like |
| AT2G24120 | SCA3 | DNA/RNA polymerases superfamily protein |

Table 1. continued
| Gene ID | Name | Description |
| --- | --- | --- |
| AT3G09650 | HCF152 | Tetratricopeptide repeat (TPR)-like superfamily protein |
| AT1G54575 | AT1G54575 | protein_coding_gene |
| AT1G66540 | AT1G66540 | Cytochrome P450 superfamily protein |
| AT5G25120 | CYP71B11 | Cytochrome p450, family 71, subfamily B, polypeptide 11 |
| AT1G56300 | AT1G56300 | Chaperone DnaJ-domain superfamily protein |
| AT1G78600 | LZF1 | light-regulated zinc finger protein 1 |
| AT3G21560 | UGT84A2 | UDP-Glycosyltransferase superfamily protein |
| AT1G75370 | AT1G75370 | Sec14p-like phosphatidylinositol transfer family protein |
| AT3G23870 | AT3G23870 | Protein of unknown function (DUF803) |
| AT1G33260 | AT1G33260 | Protein kinase superfamily protein |
| AT2G32390 | GLR3.5 | glutamate receptor 3.5 |
| AT5G19540 | AT5G19540 | protein_coding_gene |
| AT5G02860 | AT5G02860 | Pentatricopeptide repeat (PPR) superfamily protein |
| AT5G03240 | UBQ3 | polyubiquitin 3 |
| AT2G40840 | DPE2 | disproportionating enzyme 2 |
| AT3G48000 | ALDH2B4 | aldehyde dehydrogenase 2B4 |
| AT3G01310 | AT3G01310 | Phosphoglycerate mutase-like family protein |
| AT1G10760 | SEX1 | Pyruvate phosphate dikinase, PEP/pyruvate binding domain |

**Table 2.** The list of 18 genes of which decreased expression in *ref2* and *ref5* is restored by the disruption of MED5

|  |  |  |
| --- | --- | --- |
| AT2G27402 | AT2G27402 | protein_coding_gene |
| AT5G22500 | FAR1 | fatty acid reductase 1 |
| AT1G01060 | LHY | Homeodomain-like superfamily protein |
| AT3G22540 | AT3G22540 | Protein of unknown function (DUF1677) |
| AT1G74670 | GASA6 | Gibberellin-regulated family protein |
| AT1G11670 | AT1G11670 | MATE efflux family protein |
| AT2G27060 | AT2G27060 | Leucine-rich repeat protein kinase family protein |
| AT1G25230 | AT1G25230 | Calcineurin-like metallo-phosphoesterase superfamily protein |
| AT2G32720 | CB5-B | cytochrome B5 isoform B |
| AT1G65930 | cICDH | cytosolic NADP -dependent isocitrate dehydrogenase |
| AT2G40610 | EXPA8 | expansin A8 |
| AT4G13840 | AT4G13840 | HXXXD-type acyl-transferase family protein |
| AT5G48930 | HCT | hydroxycinnamoyl-CoA shikimate/quinate hydroxycinnamoyl transferase |
| AT1G06080 | ADS1 | delta 9 desaturase 1 |
| AT2G38120 | AUX1 | Transmembrane amino acid transporter family protein |
| AT1G78090 | TPPB | trehalose-6-phosphate phosphatase |
| AT1G24260 | SEPALLATA3 | K-box region and MADS-box transcription factor family protein |
| AT5G16570 | GLN1;4 | glutamine synthetase 1;4 |

**Figure 2.**
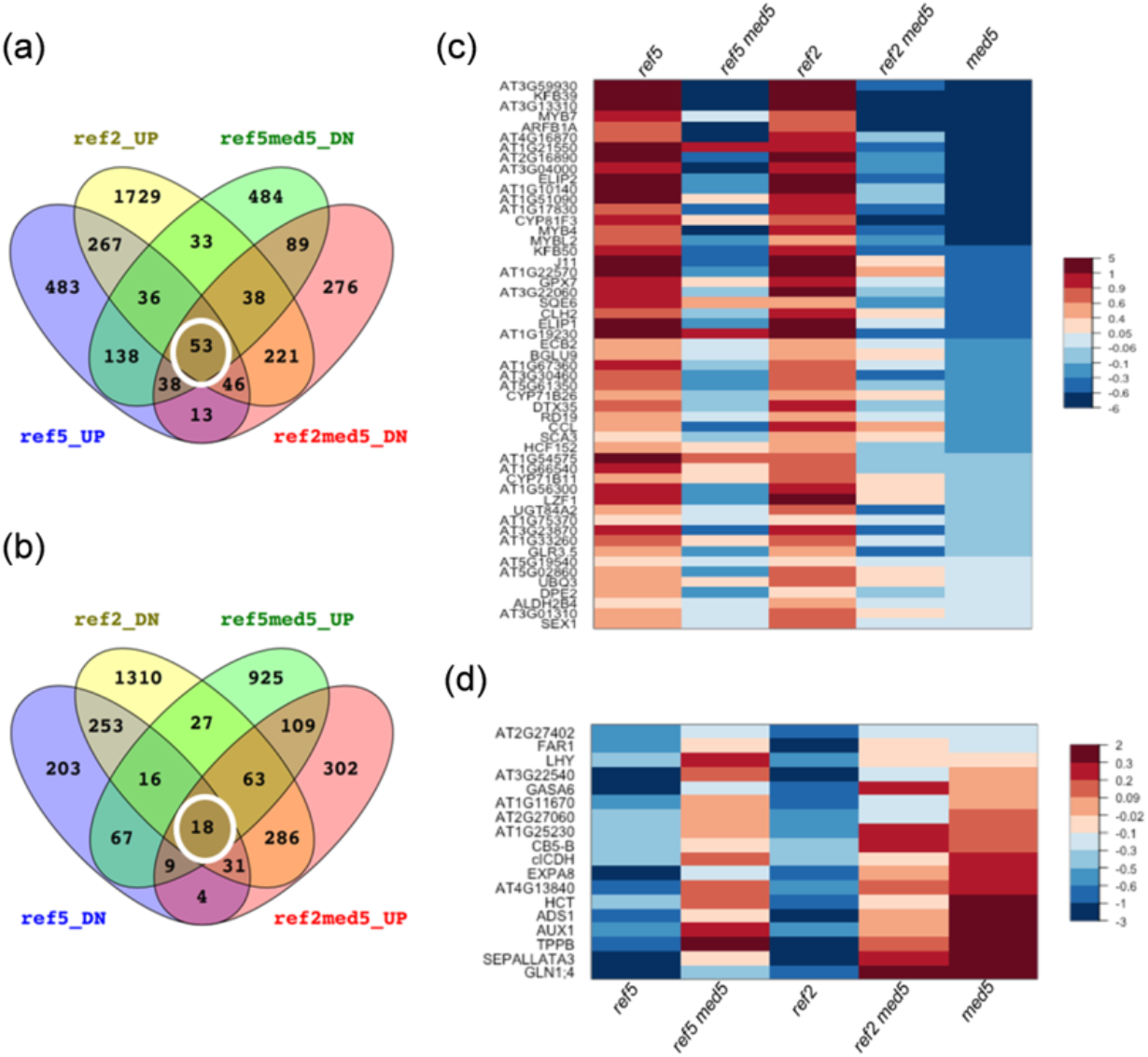
RNAseq analysis identified 71 genes that are differentially expressed in *ref5* and *ref2* mutants and of which altered expression is restored by the disruption of MED5. (a) Shown are the numbers of genes that are up-regulated in *ref5* (ref5_UP) and *ref2* (ref2_UP) compared to wild type or down-regulated in *ref5 med5* (ref5med5_DN) and *ref2 med5* (ref2med5_DN) compared to *ref5* and *ref2* respectively. (b) Number of genes that are down-regulated in *ref5* (ref5_DN) and *ref2* (ref2_DN) compared to wild type or up-regulated in *ref5 med5* (ref5med5_UP) and *ref2 med5* (ref2med5_UP) compared to *ref5* and *ref2* respectively. (c) Heat map with 53 genes that show increased expression in *ref5* and *ref2* but not in *ref5 med5* and *ref2 med5*. Color indicates log_2_ fold changes of expression in mutants compared to wild type. (d) Heat map with 18 genes that show decreased expression in *ref5* and *ref2* but not in *ref5 med5* and *ref2 med5*. Color indicates log_2_ fold changes of expression in mutants compared to wild type.

### Altered expression of *MYB4*, *BRT1* and *HCT* does not play a major role in glucosinolate/phenylpropanoid crosstalk

Among the 71 candidate genes, two phenylpropanoid biosynthetic genes, *hydroxycinnamoyl-CoA shikimate/quinate hydroxycinnamoyl transferase* (*HCT*) and *Bright Trichomes 1* (*BRT1*) and one phenylpropanoid-related transcription factor, MYB4, were identified (Table 1, 2, Fig. 3a-c). The expression of *HCT* was decreased whereas the expression of *BRT1* was increased in *ref5* and *ref2* (Fig. 3a-b). *BRT1* (At3g21560) encodes UGT84A2, a glucosyltransferase previously shown to have activity toward sinapic acid as a substrate (Sinlapadech *et al.*, 2007). Like the *ref5* and *ref2* mutants, the *brt1* mutant accumulates less sinapoylmalate than wild type. Since the expression of *BRT1* was increased in both *ref5* and *ref2*, it seemed unlikely that the enhanced expression of *BRT1* is directly involved in phenylpropanoid repression in *ref5* and *ref2* (Fig. 2c, d, Fig. 3a).

**Figure 3.**
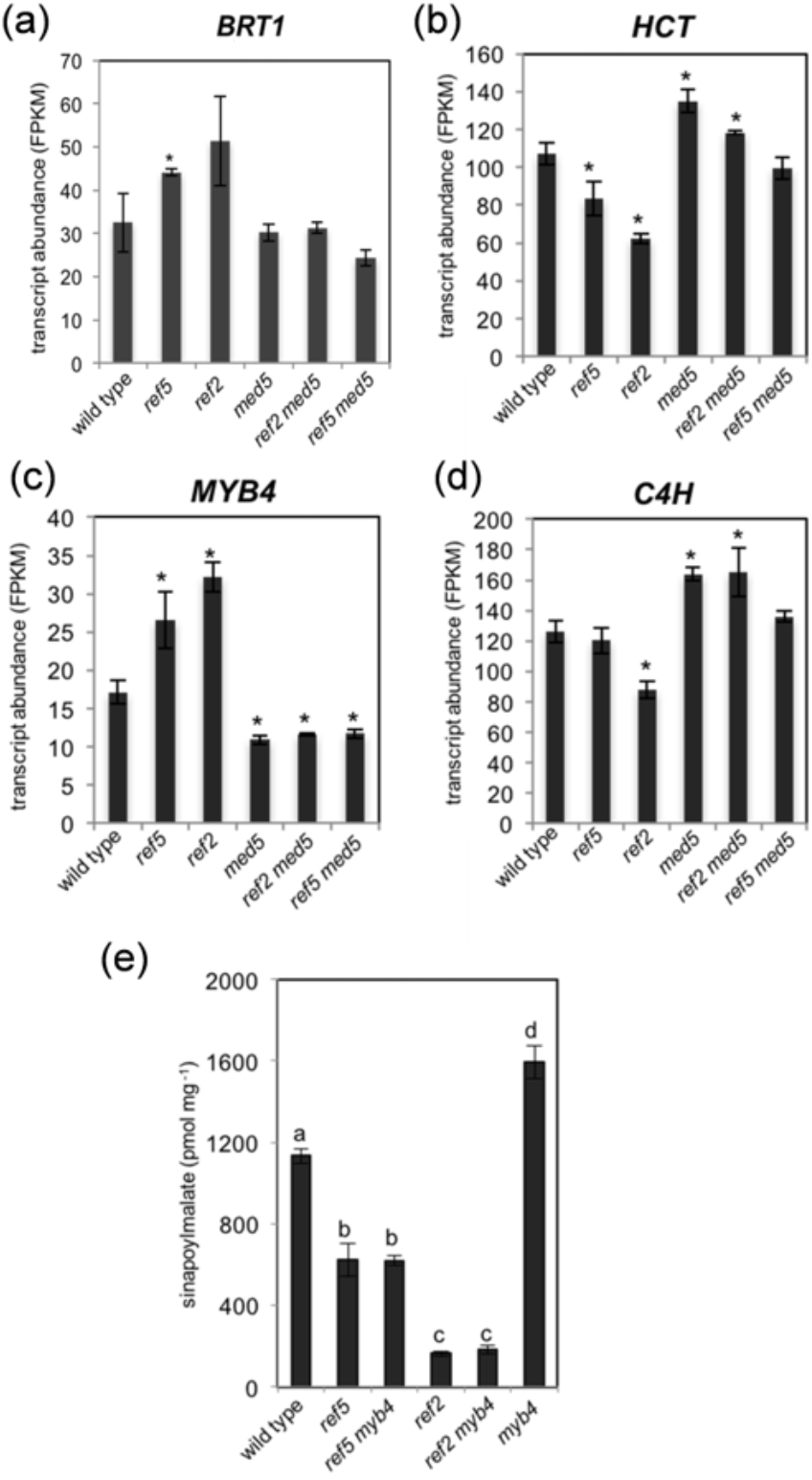
Alteration of MYB4, BRT1 or HCT does not increase phenylpropanoid production in *ref5* and *ref2*. (a-d) Expression of *BRT1*, *HCT, MYB4*, and *C4H* is shown. Transcript abundance was calculated from RNAseq data. FPKM stands for Fragments Per Kilobase of transcript per Million mapped reads. Data represent mean ± SD (n=3). ‘*’ represents the significant difference of gene expression compared to wild type (*p* < 0.05, Student t-test). (e) Sinapoylmalate content in leaves of wild type, *ref5*, *ref5 myb4, ref2, ref2 myb4*, and *myb4* mutants. Data represent mean ± SD (n=4). The means were compared by one-way ANOVA, and statistically significant differences (*p* < 0.05) were identified by Tukey’s test and indicated by a to d to represent difference between groups.

HCT catalyzes the synthesis of *p*-coumaroyl shikimate in the lignin biosynthetic pathway (Hoffmann *et al.*, 2004; Li *et al.*, 2010). The expression of *HCT* was reduced by about 15% and 30% in *ref5* and *ref2* mutants, respectively (Fig. 3b). We also excluded this gene from further consideration because previous studies with *HCT* RNAi lines showed that obvious metabolic phenotypes were not observed until HCT activity was reduced to less than 2% of the wild-type value (Hoffmann *et al.*, 2004; Li *et al.*, 2010). In addition, the repression of *HCT* results in the accumulation of flavonoids but *ref5* and *ref2* mutants accumulate less flavonoids than wild type (Fig. S1) (Hoffmann *et al.*, 2004; Li *et al.*, 2010; Kim *et al.*, 2015).

MYB4 is a transcriptional factor the function of which includes the repression of *C4H* expression(Jin *et al.*, 2000). *C4H* expression was reduced about 20% in *ref2* but it was not altered in *ref5* compared to wild type (Fig. 3d). To further evaluate the possible impact of altered *MYB4* expression, we generated *ref5 myb4* and *ref2 myb4* double mutants. In agreement with the function of MYB4, *myb4* mutants produced more sinapoylmalate than wild type (Fig. 3e) (Jin *et al.*, 2000; Panda *et al.*, unpublished) but the level of sinapoylmalate in *ref5 myb4* and *ref2 myb4* mutants was not different from that in *ref5* and *ref2* mutants (Fig. 3e). Taken together, we concluded that the altered expression of *MYB4* is not related to phenylpropanoid repression in *ref5* and *ref2*.

### The expression of *KFB*s functioning in PAL degradation is affected in *ref5* and *ref2*

Besides *HCT, BRT1* and *MYB4*, two additional genes, At3g59930 and At2g44130, caught our attention due to their known functions in phenylpropanoid metabolism. At3g59930 and At2g44130 encode the kelch-domain containing F-box (KFB) proteins KFB39 and KFB50, respectively, both of which are involved in the ubiquitination and proteasome-mediated degradation of PAL (Zhang *et al.*, 2015; Zhang *et al.*, 2013). As shown in Fig. 4, the expression of *KFB39* and *KFB50* in both *ref5* and *ref2* mutants was increased two- to four-fold compared to wild type and restored in *ref5 med5* and *ref2 med5*. It is notable that the expression of *KFB39* and *KFB50* was significantly reduced in *med5* compared to wild type (Fig. 4a, b) indicating that MED5 is required for the activation of *KFB39* and *KFB50* expression not only in *ref5* and *ref2* mutants but also in wild type. Two other KFBs, KFB1 and KFB20, function redundantly with KFB39 and KFB50 in PAL ubiquitination (Zhang *et al.*, 2013). We found that the expression of *KFB1* and *KFB20* were generally increased in *ref5* and *ref2* mutants compared to wild type but not returned to normal in *ref5 med5* and *ref2 med5* (Fig. 4c, d). These data suggest that there may be MED5-dependent and MED5-independent mechanisms for the regulation of KFB expression.

**Figure 4.**
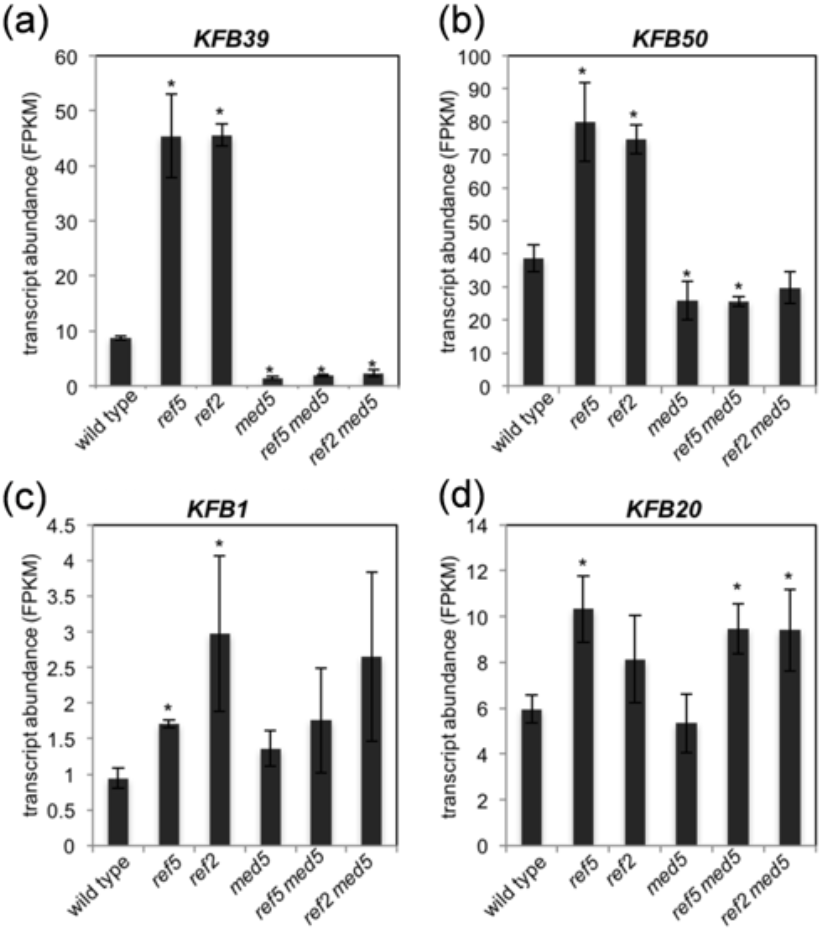
*KFB1*, *KFB20*, *KFB39* and *KFB50* are up-regulated in *ref5* and *ref2*. (a, b) Expression of *KFB39* and *KFB50* is increased in *ref5* and *ref2* mutants but not in *ref5 med5* and *ref2 med5*. (c, d) Expression of *KFB1* and *KFB20* is increased in *ref5* and *ref2* mutants but the altered expression was not restored by disruption of MED5. Transcript abundance was calculated from RNAseq data. FPKM stands for Fragments Per Kilobase of transcript per Million mapped reads. Data represent mean ± SD (n=3). * indicates *p* < 0.05 (Student’s *t*-test) when compared with wild type.

**Figure 5.**
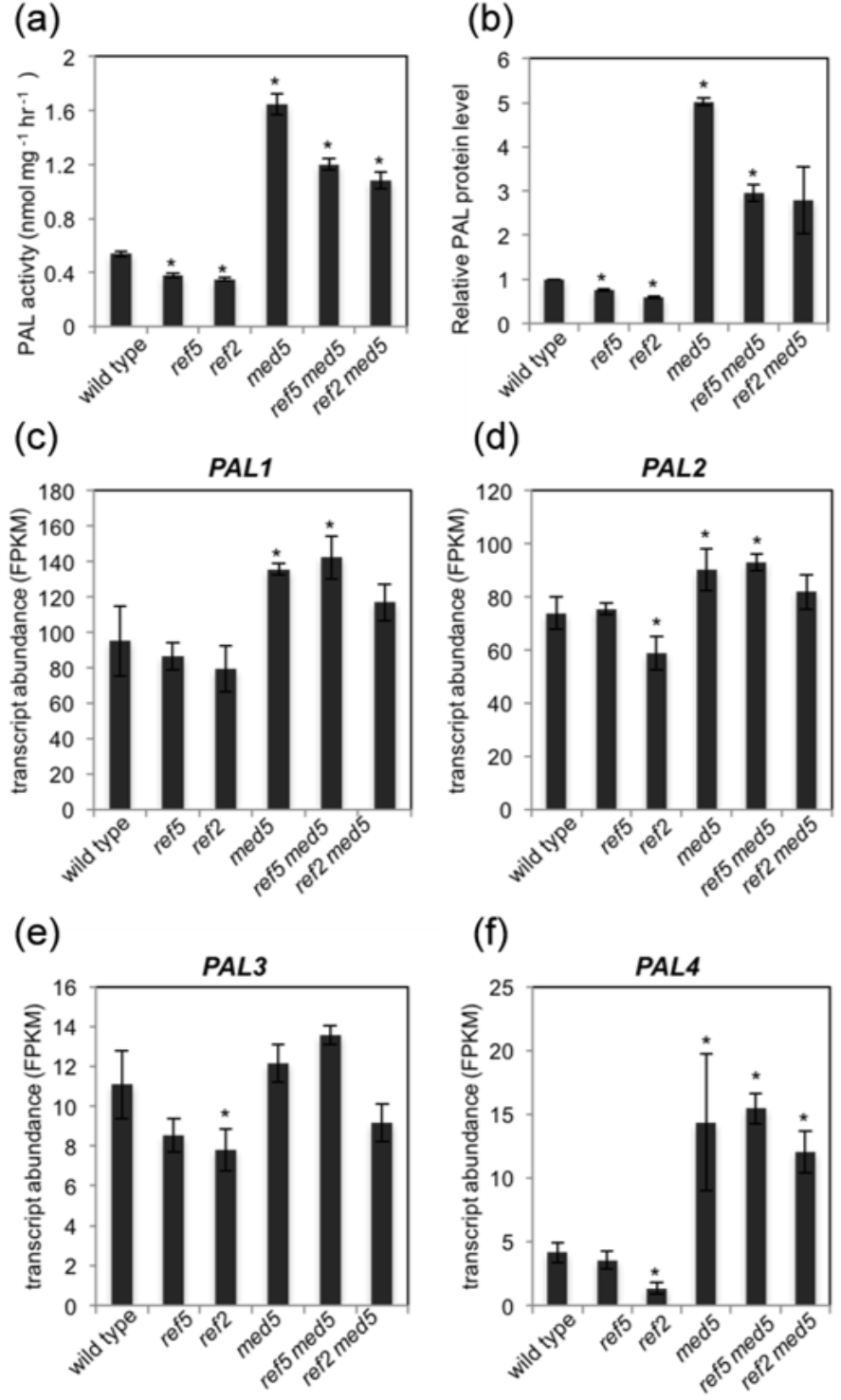
PAL activity and the amount of PAL protein are reduced in both *ref5* and *ref2* whereas the expression of PAL is not substantially affected in *ref5*. (a, b) PAL activity and relative PAL protein levels in crude extracts of whole rosette leaves from three-week-old plants are shown. PAL activity is expressed as average PAL activity in mg^−1^ protein ± SD (n=3). ‘*’ indicate *p* < 0.05 compared to wild type. *p*-values are from Student’s t-test compared to wild type. (c-f) Transcript abundance of *PAL1-PAL4* was calculated from RNAseq data. FPKM stands for Fragments Per Kiloase of transcript per Million mapped reads. Data represent mean ± SD (n=3). * indicates *p* < 0.05 (Student’s *t*-test) when compared with wild type.

In addition to the role of these four KFBs in phenylpropanoid metabolism, KFB^CHS^ is involved in the ubiquitination of chalcone synthase (CHS), an enzyme required for flavonoid production (Zhang *et al.*, 2017). We found that although *ref5* and *ref2* mutants showed reduced flavonoid content (Fig. S1), the expression of *KFB*^*CHS*^ was not affected (Fig. S2). This suggests that aldoxime-induced misregulation of KFBs is restricted to specific targets in these mutants.

### PAL activity and the amount of PAL protein are decreased in *ref5* and *ref2* mutants

Because the four KFBs we identified play a role in PAL degradation, we hypothesized that *ref5* and *ref2* mutants have enhanced PAL turnover, leading to the repression of PAL activity and phenylpropanoid production. Indeed, we found that PAL activity and protein levels were slightly reduced in *ref5* and *ref2* mutants and enhanced by elimination of MED5 (Fig 5a, b) but PAL protein and enzyme levels were lower in *ref2 med5* and *ref5 med5* than in *med5* alone, again indicating that MED5-independent mechanisms may also play a role in the crosstalk. Although PAL expression is increased in the *med5* mutant (Bonawitz *et al.*, 2014) (Fig 5c-f), it was affected only modestly in the *ref2* and *ref5* mutants suggesting that reduced PAL activity and PAL protein level in *ref2* and *ref5* (Fig 5a-b) is largely due to KFB-dependent PAL degradation. To evaluate this hypothesis, we generated multiple mutants combining *ref2* and *ref5* with KFB knockout lines. Elimination of all four KFB genes led to a substantial increase in PAL activity that was unaffected by the introduction of the *ref2* or *ref5* mutations into the same genetic background (Fig 6) indicating that the reduction of PAL activity in these mutants is due to KFB-mediated protein turnover. In contrast, PAL activity in *ref5 kfb1/20/50* or *ref2 kfb1/20/50* quadruple mutants was significantly lower than that in *kfb1/20/50* triple mutant suggesting that the accumulation of aldoximes in *ref2* and *ref5* affects PAL activity through KFB39 (Fig 6).

**Figure 6.**
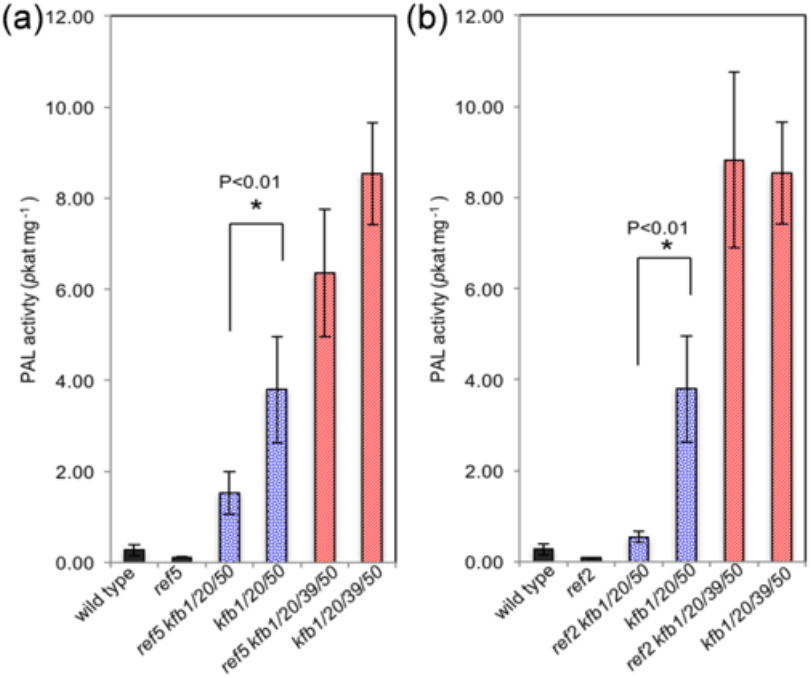
Disruption of KFBs restores PAL activity in *ref5* and *ref2*. (a-b) PAL activity in *ref5* (a) and *ref2* (b) when KFBs were disrupted. PAL activity was measured in crude extracts of rosette leaves from three-week-old plants. PAL activity is expressed as average PAL activity in mg^−1^ protein ± SD (n=3). Blue circle bars show PAL activity when KFB1, KFB20 and KFB50 were disrupted in *ref5* or *ref2* compared to *kfb1 kfb20 kfb50* triple mutant. Red diagonal bars show PAL activity when all four KFBs (KFB1, KFB20, KFB39 and KFB50) were disrupted in *ref5* or *ref2* compared to *kfb1 kfb20 kfb39 kfb50* quadruple mutant. Data represent mean ± SD (n=3). *p*-values are from Student’s *t*-test compared to two mutants indicated with solid lines.

### KFB-mediated PAL degradation is one of multiple mechanisms involved in the repression of phenylpropanoid production in *ref5* and *ref2*

Given that disruption of KFBs restored PAL activity to *ref2* and *ref5*, we tested whether this restoration would rescue the reduced phenylpropanoid production in *ref5* and *ref2* (Fig. 7). Because the expression of only *KFB39* and *KFB50* is increased in *ref5* and *ref2* in a MED5-dependent manner (Fig.4), we first tested the impact of these two genes on phenylpropanoid production in *ref5*. The knockout of either KFB39 or KFB50 alone did not affect sinapoylmalate accumulation in *ref5* mutant (Fig. 7a). When both KFB39 and KFB50 were eliminated, sinapoylmalate content was slightly increased compared to the *ref5* mutant but its level was still less than wild type (Fig. 7a). When all four of the KFBs were removed, sinapoylmalate content in the *ref5* and *ref2* mutant backgrounds was restored to the wild-type level (Fig. 7b) but was not as high as that in *kfb1/20/39/50* quadruple mutant (Fig. 7b). Considering that there is no statistical significance among PAL activities of these mutants (Fig. 6a, b), in addition to PAL, there must be other step(s) of the phenylpropanoid pathway that are affected in *ref5* and *ref2* mutants in addition to PAL.

**Figure 7.**
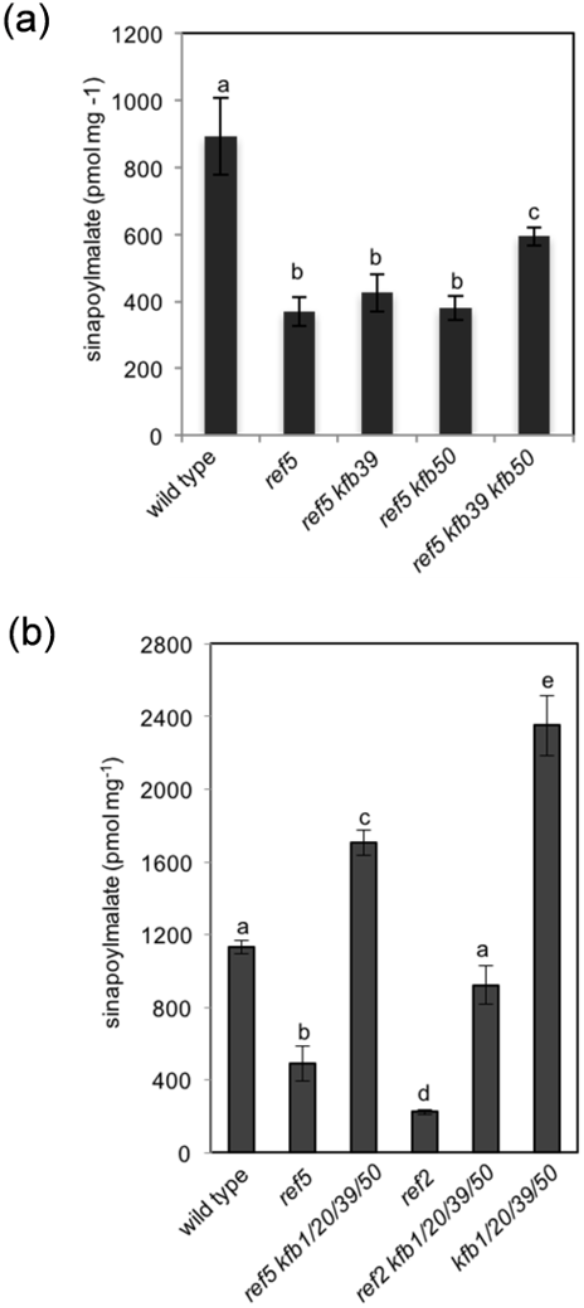
Disruption of KFBs restores phenylpropanoid content in *ref5* and *ref2*. (a) Sinapoylmalate content in leaves of wild type, *ref5*, *ref5 kfb39, ref5 kfb50*, and *ref5 kfb39 kfb50* mutants. Data represent mean ± SD (n=4). The means were compared by one-way ANOVA, and statistically significant differences (*p* < 0.05) were identified by Tukey’s test and indicated by a to c to represent difference between groups. (b) Sinapoylmalate content in *ref5* and *ref2* mutants when KFBs were disrupted. Data represent mean ± SD (n=3). The means were compared by one-way ANOVA, and statistically significant differences (*p* < 0.05) were identified by Tukey’s test and indicated by a to e to represent difference between groups.

**Figure 8.**
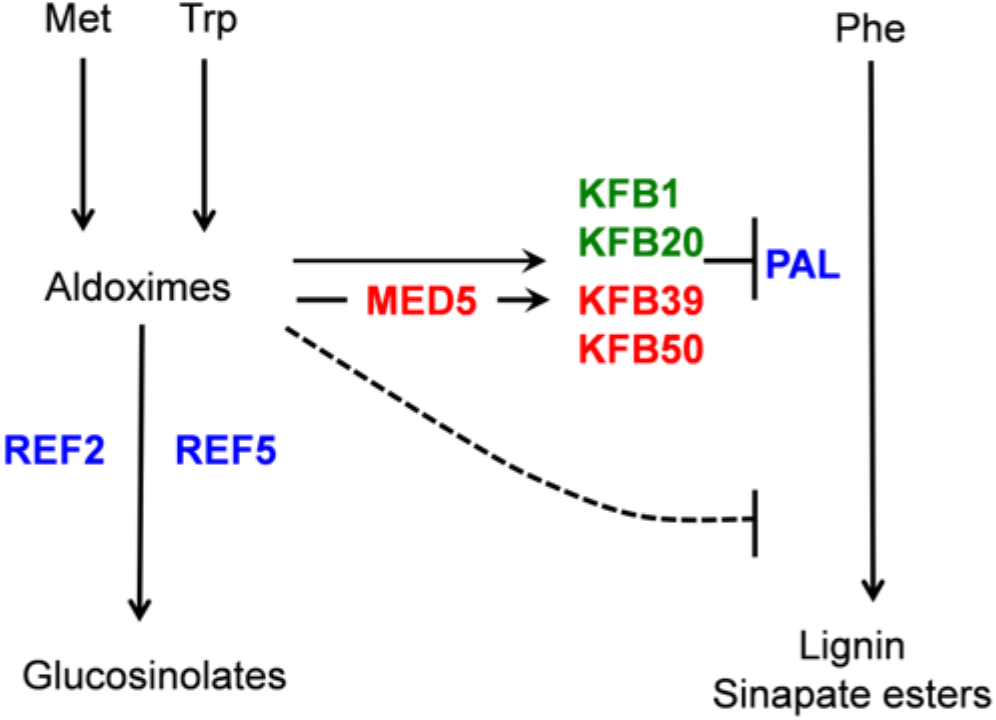
Model for the crosstalk between glucosinolate biosynthesis and phenylpropanoid biosynthesis. Accumulation of aldoximes or their derivatives activate the expression of *KFB1*, *KFB20*, *KFB39* and *KFB50*, which affects the degradation of PAL functioning at the entry point of phenylpropanoid biosynthesis pathway. Besides PAL, our data suggest that additional step(s) in phenylpropanoid pathway are affected by altered level of aldoximes or their derivatives.

## Discussion

Glucosinolates and phenylpropanoids are secondary metabolites that are made through unique biosynthetic pathways. The first step of glucosinolate core structure formation is the production of aldoximes, which are subsequently used as substrates by CYP83A1/REF2 or CYP83B1/REF5 in *Arabidopsis*. Previous studies with two *Arabidopsis* mutants have demonstrated an unexpected link between glucosinolate and phenylpropanoid production (Hemm *et al.*, 2003; Kim *et al.*, 2015). It appears that the accumulation of aldoximes or their derivatives limits phenylpropanoid production (Hemm *et al.*, 2003; Kim *et al.*, 2015). Because aldoximes are precursors of many other biomolecules besides glucosinolates, including cyanogenic glucosides and nitriles, and can be found in diverse plant species such as maize, poplar, yew and coca (Irmisch *et al.*, 2013, 2015; Luck *et al.*, 2017; Sørensen *et al.*, 2018; Bhalla *et al.*, 2018; Dhandapani *et al.*, 2019), aldoxime-mediated suppression of phenylpropanoid metabolism may be relevant to other species and metabolic pathways beyond Arabidopsis.

Previous work showed that glucosinolate/phenylpropanoid crosstalk requires MED5 (Kim *et al.*, 2015) but its underlying mechanism was not identified. In this study, we demonstrate that the crosstalk involves transcriptional regulation of four *KFB* genes functioning in PAL ubiquitination and that MED5 is required for the upregulation of two of them. Interestingly, MED5 is only required for the increased expression of *KFB39* and *KFB50* in the context of the *ref5* and *ref2* mutations, but not for the expression of *KFB1* and *KFB20* (Fig 4). Apparently, there exist distinct regulatory mechanisms between the two sub families of KFB proteins even though they function redundantly in PAL degradation. In contrast, the expression of *KFB*^*CHS*^, which functions in flavonoid biosynthesis, (Zhang *et al.*, 2017) was not affected in the high aldoxime containing mutants, although both showed reduced flavonoid contents indicating that aldoxime metabolism affects a specific group of KFBs and not others (Kim *et al.*, 2015) (Fig. S1, S2). Instead, KFB-mediated PAL degradation and the ensuing reduction in PAL activity is a more likely explanation for the reduced flavonoid content of *ref5* and *ref2*.

We previously suspected that the influence of aldoxime metabolites on phenylpropanoid metabolism might be just a pharmacological effect rather than a biologically relevant regulatory mechanism; however, the data presented here suggest that there may be a more sophisticated function for this crosstalk. First, at least one aspect of the crosstalk requires a specific subunit of the transcriptional regulatory complex Mediator (Kim *et al.*, 2015) which enhances the expression of KFB39 and KFB50 and thereby reduces PAL activity (Fig. 4a, b). Second, this regulatory circuit affects a certain group of KFB genes but not others, indicating some level of specificity to the process (Fig. 4, Fig. S2). Finally, a certain level of aldoxime-sensitivity is maintained in KFB-deficient lines, indicating that disruptions in glucosinolate metabolism impinge on phenylpropanoid metabolism in multiple ways. Further study will be required to elucidate how plants sense aldoximes or their derivatives and transduce signals to regulate gene expression and enzyme activity.

The phenylpropanoid pathway is subject to multiple layers of regulation at various steps. As PAL is generally considered to be the first rate-limiting enzyme in the phenylpropanoid pathway, the transcriptional and post-translational modification regulation of its activity can modulate the production of a variety of phenolic compounds. For example, developmental cues and environmental stresses affect PAL expression and its activity is highly correlated with PAL transcript abundance (Olsen *et al.*, 2008; Huang *et al.*, 2010; Zhang *et al.*, 2013). Further, ubiquitination or phosphorylation of PAL can affect its stability or specific subcellular localization (Zhang *et al.*, 2013, 2015; Allwood *et al.*, 1999; Cheng *et al.*, 2001). Finally, PAL is subject to feedback regulation through biosynthetic intermediates (Blount *et al.*, 2000; Bolwell *et al.*, 1986). Cinnamic acid, a product of PAL can influence its expression and activity (Mavandad *et al.*, 1990; Blount *et al.*, 2000) and various flavonoids including naringenin, a precursor of flavonoids, can inhibit its activity (Sato & Sankawa, 1983; Yin *et al.*, 2012). Our data now suggest that an intermediate in glucosinolate biosynthesis or a derivative thereof, made through an independent pathway, influences PAL activity by regulating its protein stability.

The fact that KFBs are at least partially responsible for the phenylpropanoid deficiency in *ref5* and *ref2*, and that REF5 and REF2 act in parallel glucosinolate pathways suggests that there may be a common metabolite or class of metabolite that triggers the crosstalk in these mutants. Although both CYP83A1/REF2 and CYP83B1/REF5 have activity toward various aldoximes, they show distinctive substrate specificity (Naur *et al.*, 2003). CYP83A1/REF2 mainly metabolizes the aliphatic oximes derived from chain-elongated homologs of methionine. In contrast, CYP83B1/REF5 has a higher affinity for indole-3-acetaldoxime (IAOx), with a 50-fold difference in K_m_ between CYP83A1/REF2 and CYP83B1/REF5 for IAOx (Naur *et al.*, 2003). Consistently, plants having defects in CYP83B1/REF5 exhibit characteristic phenotypes that are associated with increased level of IAOx-derived auxin (Boerjan *et al.*, 1995; Wagner *et al.*, 1997; Barlier *et al.*, 2000; Bak & Feyereisen, 2001; Kim *et al.*, 2015), whereas the *ref2* mutant does not (Hemm *et al.*, 2003). Thus, it seems improbable that the level of IAOx or its derivatives is increased in *ref2*. Interestingly, *Arabidopsis* transgenic plants overexpressing sorghum *CYP79A1*, or both *CYP79A1* and *CYP71E1* together showed reduced production of phenylpropanoids such as sinapoylglucose, sinapoylmalate and flavonoids (Kristensen *et al.*, 2005). Since sorghum CYP79A1 has specific activity toward tyrosine to produce *p*-hydroxyphenylacetaldoxime (Kristensen *et al.*, 2005), the accumulation of *p*-hydroxyphenylacetaldoxime or its derivatives may also affect phenylpropanoid production. Given that the altered level of various types of aldoxime results in similar impact on phenylpropanoid production, it is possible that a common structure of the oximes, rather than the precise identity of the amino acid R-group from which the compound was derived may be the factor important for the crosstalk.

A recent study identified scaffolding proteins that structure the monolignol biosynthetic P450 enzyme complex (monolignol metabolon) in the endoplasmic reticulum membrane (Gou *et al.*, 2018). This complex appears to function in regulating monolignol biosynthesis but not flavonoid synthesis (Gou *et al.*, 2018). It has been proposed that metabolic complexes of dhurrin and glucosinolate biosynthetic enzymes play a role in optimizing the production of these phytochemicals with the CYP79 aldoxime-producing enzymes and the CYP71E1 or CYP83 aldoxime-consuming enzymes forming similar complexes on the endoplasmic reticulum membrane (Møller, 2010). Considering that aldoximes or their derivatives can influence phenylpropanoid production by affecting PAL, the evolution of these complexes may have been important not only to increase local concentration of metabolites but also to reduce leakage of biologically active metabolites that can affect other pathways.

In summary, our transcriptome analyses and genetic study with a set of glucosinolate deficient mutants reveal that the accumulation of aldoximes or their derivatives affects expression of a specific group of F-box genes, thereby altering the stability of PAL as well as act to negatively affect at least one other step of phenylpropanoid metabolism. Considering that aldoximes are precursors of important biomolecules and the oxime-producing CYP79s are widespread in higher plants (Sørensen *et al.*, 2018), this crosstalk may be relevant to not only glucosinolates but also other metabolic pathways.

## ACKNOWLEDGMENTS

The authors thank to Dr. Jian-Kang Zhu for generous gift with pCrisper-CAS9 vector and Dr. Xu (Sirius) Li for *myb4* mutant seeds. This work was supported as part of the Center for Direct Catalytic Conversion of Biomass to Biofuels, an Energy Frontier Research Center funded by the U.S. Department of Energy, Office of Science, Basic Energy Sciences under Award # DE-SC0000997.

## Author contributions

J.K., and C.C. conceived and designed the experiments; J.K, and X.Z performed the experiments; J.K, X.Z, P.P., C-J.L, and C.C., analyzed and interpreted the data; J.K. and C.C. wrote the manuscript;

## Data availability

The authors declare that all data supporting the findings of this study are included in the manuscript and Supplementary Information files are available from the corresponding authors upon request. RNAseq data have been deposited in NCBI’s Gene Expression Omnibus^60^ under GEO Series accession number GSE99581.

## Supplementary information

The following materials are available in the online version of this article.

**Supplemental Figure S1.** Flavonoid content was not increased in *ref5* and *ref2* compared to wild type or in *ref5 kfb1/20/39/50* and *ref2 kfb1/20/39/50* compared to *kfb1/20/39/50*.

**Supplemental Figure S2**. Expression of *KFB*^*CHS*^ was not significantly altered in *ref5* and *ref2*.

