## Supplemental File for "Glucosinolate and phenylpropanoid biosynthesis are linked by proteasome-dependent degradation of PAL"

Article acceptance date: Click here to enter a date.

The following Supporting Information is available for this article:

**Fig. S1** Flavonoid content was not increased in *ref5* and *ref2* compared to wild type or in *ref5 kfb1/20/39/50* and *ref2 kfb1/20/39/50* compared to *kfb1/20/39/50*.

**Fig. S2** Expression of $KFB^{CHS}$ was not significantly altered in *ref5* and *ref2*.

**Fig. S1** Flavonoid content was not increased in *ref5* and *ref2* compared to wild type or in *ref5 kfb1/20/39/50* and *ref2 kfb1/20/39/50* compared to *kfb1/20/39/50*. K1, kaempferol 3-*O*-[6”-*O*-(rhamnosyl) glucoside] 7-*O*-rhamnoside (a); K2, kaempferol 3-*O*-glucoside 7-*O*-rhamnoside (b) ; K3, kaempferol 3-*O*-rhamnoside 7-*O*-rhamnoside (c) are shown. ‘*’ indicate $p < 0.05$ compared to wild type. *p*-values are from Student’s t-test compared to wild type (n=4).

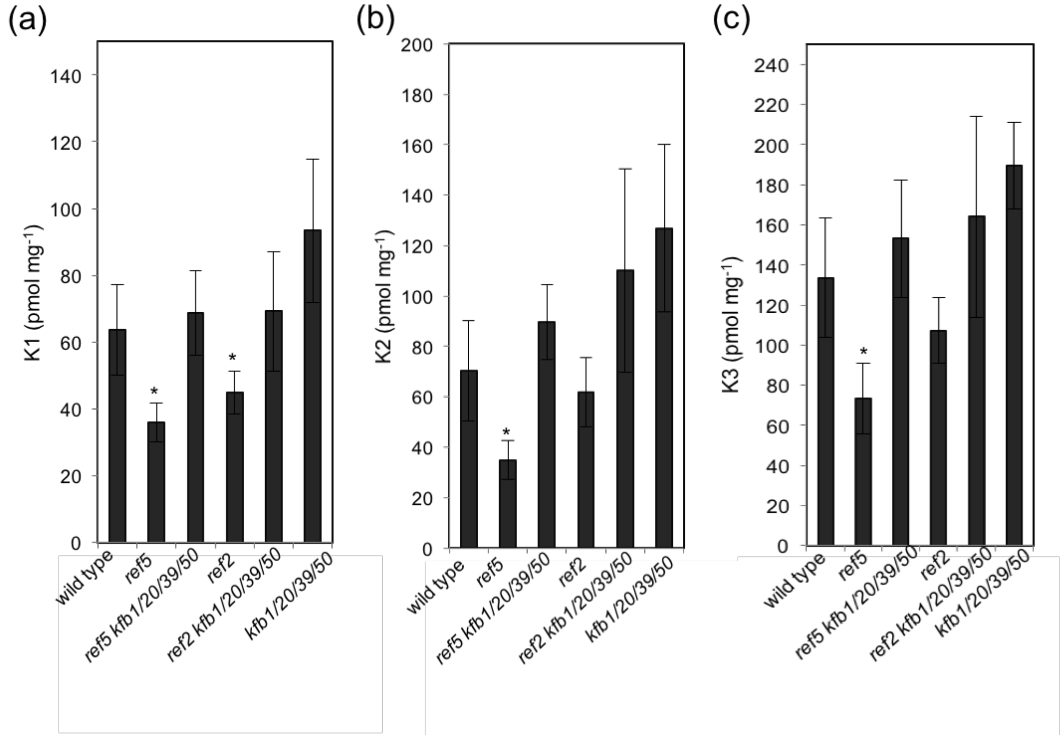


**Fig. S2** Expression of $KFB^{CHS}$ was not significantly altered in *ref5* and *ref2*. FPKM stands

for Fragments Per Kilobase of transcript per Million mapped reads. Data represent mean ± SD (n=3).

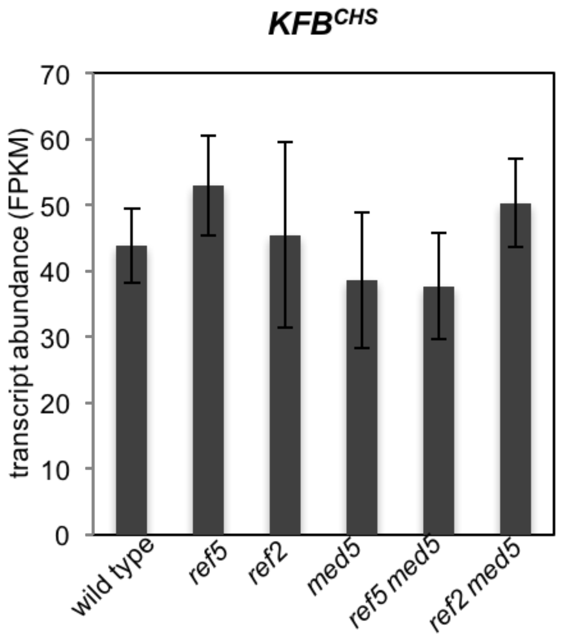
